# Sex differences in the adult human brain: Evidence from 5,216 UK Biobank participants

**DOI:** 10.1101/123729

**Authors:** Stuart J. Ritchie, Simon R. Cox, Xueyi Shen, Michael V. Lombardo, Lianne M. Reus, Clara Alloza, Mathew A. Harris, Helen L. Alderson, Stuart Hunter, Emma Neilson, David C. M. Liewald, Bonnie Auyeung, Heather C. Whalley, Stephen M. Lawrie, Catharine R. Gale, Mark E. Bastin, Andrew M. McIntosh, Ian J. Deary

## Abstract

Sex differences in the human brain are of interest, for example because of sex differences in the observed prevalence of psychiatric disorders and in some psychological traits. We report the largest single-sample study of structural and functional sex differences in the human brain (2,750 female, 2,466 male participants; 44-77 years). Males had higher volumes, surface areas, and white matter fractional anisotropy; females had thicker cortices and higher white matter tract complexity. There was considerable distributional overlap between the sexes. Subregional differences were not fully attributable to differences in total volume or height. There was generally greater male variance across structural measures. Functional connectome organization showed stronger connectivity for males in unimodal sensorimotor cortices, and stronger connectivity for females in the default mode network. This large-scale study provides a foundation for attempts to understand the causes and consequences of sex differences in adult brain structure and function.

## Introduction

Sex differences have been of enduring biological interest (Darwin, 1871), but our knowledge about their relevance to the human brain is surprisingly sparse. It has been noted by several researchers that the potential influences of sex are under-explored in neuroscientific research (Beery & Zucker, 2011; Cahill, 2006, 2017; Karp, 2017). A fuller understanding of morphological and functional differences between the brains of the human sexes might provide insight into why the observed prevalence of some psychiatric disorders differs substantially by sex (Rutter et al., 2003), and would assist in explaining several behavioural sex differences (Gur & Gur, 2017; Zell et al., 2015). As biomedical research moves closer to the ideals of precision medicine (e.g. Collins & Varmus, 2015), it is even more pressing that we have a more nuanced understanding of similarities and differences in brain structure and function across the sexes. Here, we report a study that characterises multimodal sex differences in the human brain in the largest sample to date.

It is of particular importance to gain a more detailed picture of how the brains of males and females differ, because several psychiatric disorders and conditions differ in their prevalence between the sexes. For instance, rates of Alzheimer’s disease are higher in females than males, prompting a recent call for the prioritisation of biomedical research into sex differences in measures relevant to this disorder (Mazure & Swendsen, 2016). Females also show a higher prevalence of major depressive disorder (Rutter et al., 2003; Gobinath et al., 2017), whereas males display higher rates of disorders such as autism spectrum disorder (Baron-Cohen et al., 2011), schizophrenia (Aleman et al., 2003) and dyslexia (Arnett et al, 2017). Improving therapeutic strategies for these conditions will almost certainly require accurate quantitative estimates of where and how the sexes differ normatively.

Moreover, although many psychological sex differences are small (consistent with the “gender similarities hypothesis”; Hyde et al., 2014), some behaviours and traits do show reliable and substantial differences. For instance, performance on mental rotation tasks (Maeda & Yoon, 2013) and physical aggression (Archer, 2004) are on average higher in males, whereas self-reported interest in people versus things (Su et al., 2009) and the personality traits of neuroticism (Schmitt et al., 2008) and agreeableness (Costa et al., 2001) are on average higher in females. A full explanation of these cognitive and behavioural phenomena would benefit from a better understanding of how brain differences may mediate behavioural differences.

Our understanding of brain sex differences has been hampered by low statistical power in previous studies. Small-sample research has become a considerable concern in neuroscience research (Button et al., 2013; Nord et al., 2017), and the concern no less applies to research on sex differences. To illustrate this point, in the most recent meta-analysis of macrostructural sex differences in brain subregions (Ruigrok et al., 2014)—which revealed a complex pattern of differences, with both males and females showing larger brain volume depending on the brain substructure in question—studies that examined sex differences in specific sub-regions of interest (rather than in broad, overall measures) had a mean sample size of 130 participants (range = 28-465). Since the publication of that meta-analysis, some larger macrostructural studies have appeared, though they are either in younger participants only (Gennatas et al., 2017; Gur & Gur, 2016; Wierenga et al., 2017) or somewhat limited in the number of brain measures they report (Jäncke et al., 2015). Adult macrostructural studies with a large scale—both in terms of sample size and in terms of brain regions analysed—are required.

Beyond macrostructural measures, there may also be robust sex differences in measures of the brain’s white matter microstructure. Studies that have attempted to quantify sex differences in white matter microstructure using diffusion tensor MRI—which uses information about the movement of water molecules through the brain’s white matter tracts to produce measures such as fractional anisotropy, which has been linked to variation in cognitive and health-related traits (Sundgren et al., 2004)—are rare and, where they exist, small in sample size (Kanaan et al., 2012; Dunst et al., 2014). Newer Neurite Orientation Dispersion and Density Imaging (NODDI) approaches—which provide data on the more specific morphological structure and organisational complexity of the brain’s neurites (axons and dendrites; Zhang et al., 2012)—remain under-investigated with regard to factors such as sex, especially at large scale.

In addition to the above structural brain imaging measures, it is also of interest to investigate sex differences in brain function. Examinations of sex differences in resting-state functional connectivity—the functional measure used in the present study, which indexes the temporal relations between activation in anatomically-separate brain regions while the brain is at rest (that is, not completing any experimenter-directed task; van den Heuvel & Hulshoff Pol, 2010)—have also shown substantial differences, for example within the default mode network (where females show stronger connectivity) and within sensorimotor and visual cortices (where males show stronger connectivity; Biswal et al., 2010). As has been noted (Scheinost et al., 2015), a better characterisation of broad patterns, including sex differences, in relatively novel measures such as functional connectivity (and in the NODDI parameters described above) is of importance to establish a “baseline” upon which future studies of normal versus abnormal function can rely.

There is more to sex differences than averages: there are physical and psychological traits that tend to be more variable in males than females. The best-studied human phenotype in this context has been cognitive ability: almost universally, studies have found that males show greater variance in this trait (Deary et al., 2007; Johnson et al., 2008; Lakin et al., 2013; though see Iliescu et al., 2016). This has also been found for academic achievement test results (a potential consequence of cognitive differences; Lehre et al., 2009a, 2009b; Machin & Pekkarinen, 2008), other psychological characteristics such as personality (Borkenau et al., 2013), and a range of physical traits such as athletic performance (Olds et al., 2006), and both birth and adult weight (Lehre et al., 2009a). To our knowledge, only two prior studies have explicitly examined sex differences in the variability of brain structure (Wierenga et al., 2017; Lange et al., 1997), and no studies have done so in individuals older than 20 years. Here, we addressed this gap in the literature by testing the “greater male variability” hypothesis in the adult brain.

### The Present Study

To date, there exists no single, comprehensive, well-powered analysis of sex differences in mean and variance in the brain that covers structural, diffusion, and functional MRI measures. Here, we examine multimodal sex differences in adult human brain structural and functional organization in the largest and most definitive study to date, ensuring high levels of statistical power and reliability. We used data from UK Biobank (Allen et al., 2012), a biomedical study based in the United Kingdom. A subset of the full sample of 500,000 participants have contributed neuroimaging data (Miller et al., 2016); a portion of these data have been released for analysis while collection is ongoing. We tested male-female differences (in mean and variance) in overall and subcortical brain volumes, mapped the magnitude of sex differences across the cortex with multiple measures (volume, surface area, and cortical thickness), and also examined sex differences in white matter microstructure derived from DT-MRI and NODDI. We tested the extent to which these structural differences (in both broad and regional measures) mediated sex variation in scores on two cognitive tests. At the functional level, we also examined large-scale organization of functional networks in the brain using resting-state fMRI functional connectivity data and data-driven network-based analyses.

## Materials and Methods

### Participants

UK Biobank (http://www.ukbiobank.ac.uk/) is a large, population-based biomedical study comprising around 500,000 participants recruited from across Great Britain (England, Scotland, and Wales) between 2006 and 2014 (Allen et al., 2012; Collins, 2012; Miller et al, 2012). After an initial visit for the gathering of medical and other information, a subset of these participants began attending for head MRI scanning. MRI data from 5,216 participants were available for the present study (mean age = 61.72 years, SD = 7.51, range = 44.23-77.12), collected at an average of around four years after the initial visit, and completed on an MRI scanner in Manchester, UK (that is, all data in this analysis were collected on the same scanner; see below for scanner details). There were 2,750 females (mean age = 61.12 years, SD = 7.42, range = 44.64-77.12) and 2,466 males (mean age = 62.39 years, SD = 7.56, range = 44.23-76.99). Further details regarding the demographics and representativeness of the sample are reported in the Supplemental Materials.

UK Biobank received ethical approval from the Research Ethics Committee (reference 11/NW/0382). The present analyses were conducted as part of UK Biobank application 10279. All participants provided informed consent to participate. Further information on the consent procedure can be found at the following URL: http://biobank.ctsu.ox.ac.uk/crystal/field.cgi?id=200.

### Brain image acquisition and processing

MRI data for all participants were acquired on a single Siemens Skyra 3T scanner, according to previously-reported procedures (Miller et al., 2016; Online Documentation: http://biobank.ctsu.ox.ac.uk/crystal/refer.cgi?id=2367; http://biobank.ctsu.ox.ac.uk/crystal/refer.cgi?id=1977). Briefly, the acquired 3D MPRAGE T1-weighted volumes were pre-processed and analysed using FSL tools (http://www.fmrib.ox.ac.uk/fsl) by the UK Biobank brain imaging team. This included a raw, de-faced T1-weighted volume, a reduced field-of-view (FoV) T1-weighted volume, and further processing, which included skull stripping, bias field correction and gross tissue segmentation using FNIRT (Andersson et al., 2001, 2007) and FAST (Zhang et al., 2001), yielding cerebrospinal fluid (CSF), grey and white matter volumes. Head size approximation was achieved through SIENEX-style analyses (Smith, 2002) which estimates the external surface of the skull. Subcortical segmentation was also conducted by the UK Biobank imaging team using FIRST (Patenaude et al., 2011) to provide the volumes of 15 structures. These data were made available to us as a downloadable dataset of Imaging Derived Phenotypes (IDPs).

#### Subregional analyses

In addition, we used the FoV-reduced T1-weighted volumes from the first release of UK Biobank MRI data to reconstruct and segment the cortical mantle using default parameters in Freesurfer v5.3 (http://surfer.nmr.mgh.harvard.edu/; Fischl & Dale, 2000; Fischl et al., 2004; Ségonne et al., 2007), according to the Desikan-Killiany atlas (Desikan et al., 2006). Following visual checking of each segmentation (including tissue identification and boundary positioning errors), the volume, thickness and surface area of all 68 cortical regions of interest (see Figure S3) were extracted for 3,875 participants. The magnitudes of sex differences across the cortical surface were visualised using the freely-available Liewald-Cox Heatmapper tool (http://www.ccace.ed.ac.uk). We also registered the vertices of each participants’ cortical model to the freesurfer average pial surface, smoothed at 20mm full width half maximum. Vertex-wise regression analyses were then conducted across each aligned cortical vertex for volume, surface area and thickness using the SurfStat MATLAB toolbox (http://www.math.mcgill.ca/keith/surfstat) for Matrix Laboratory R2014a (The MathWorks Inc., Natick, MA).

#### White matter microstructure

MRI (dMRI) acquisition are openly available from the UK Biobank website in the form of a Protocol (http://biobank.ctsu.ox.ac.uk/crystal/refer.cgi?id=2367), Brain Imaging Documentation (http://biobank.ctsu.ox.ac.uk/crystal/refer.cgi?id=1977), and in Miller et al. (2016). Following gradient distortion correction, and further correction for head movement and eddy currents, BEDPOSTx was used to model within-voxel multi-fibre tract orientation, followed by probabilistic tractography (with crossing fibre modelling) using PROBTRACKx (Behrens et al., 2003, 2007; Jbabdi et al., 2012). The AutoPtx plugin for FSL (de Groot et al., 2013) was used to map 27 major white matter tracts from which tract-average fractional anisotropy was derived. On the basis of the factor analyses described by Cox et al. (2016), we selected 22 of the white matter tracts for inclusion in the present study. Neurite orientation dispersion and density imaging (NODDI) modelling was conducted using the AMICO tool (https://github.com/daducci/AMICO; Daducci et al., 2015), and the resultant orientation dispersion (OD) maps were registered with the AutoPtx tract masks to yield an average OD value per tract. These measures were also derived by the UK Biobank imaging team and were available as imaging-derived phenotypes (IDPs). An atlas of the selected white matter tracts is provided in Figure S4.

Note that the mean sex differences in the white matter microstructural parameters studied here were already reported by Cox et al. (2016). Here, we add the analyses of variance differences, and the mediation models with diffusion properties as the mediator of the sex difference in cognitive abilities (see below).

#### Resting-state fMRI (rsfMRI)

To analyse resting-state connectivity, we used bulk data from network matrices generated by UK Biobank. As described in the Online Methods section of Miller et al. (2016), participants lay in the scanner and were instructed to “keep their eyes fixated on a crosshair, relax, and ‘think of nothing in particular’”. Data pre-processing, group-Independent Components Analysis (ICA) parcellation, and connectivity estimation were carried out by UK Biobank using FSL packages (http://biobank.ctsu.ox.ac.uk/crystal/refer.cgi?id=1977). The following preprocessing procedures were applied: motion correction using MCFLIRT (Jenkinson et al., 2002), grand-mean intensity normalisation using a single multiplicative factor, high-pass temporal filtering with a Gaussian-weighted least-squares straight line fitting (sigma was set as 50.0s), EPI unwarping using a field map scanned before data collection, gradient distortion correction (GDC) unwarping, and removal of structural artefacts using an ICA-based X-noiseifier (Beckmann & Smith, 2004). Any gross preprocessing failure was visually checked and eliminated (Miller et al., 2016). Group-ICA parcellation was conducted on 4,162 participants. The preprocessed EPI images were fed into the MELODIC tool in FSL to generate 100 distinct ICA components. The spatial maps for the components are available at the following URL: http://www.fmrib.ox.ac.uk/datasets/ukbiobank/index.html. Details of preprocessing steps can be found in pages 12, 15 and 16 of Brain Imaging Document (version 1.3) from UK Biobank data showcase website: https://biobank.ctsu.ox.ac.uk/crystal/docs/brain_mri.pdf.

Time series data from the 55 components were used for connectivity analysis, with each component as a node. Two 55×55 matrices of fully-normalized temporal correlations and partial temporal correlations were derived for each participant. A larger absolute number indicates stronger temporal connectivity, and the valence represents whether the connection is positive or negative. Partial temporal correlation matrices were used for analysis, as they represent direct connections better than full temporal correlations. Estimation of the partial correlation coefficients was conducted using FSLnets package in FSL (https://fsl.fmrib.ox.ac.uk/fsl/fslwiki/FSLNets). To produce a sparser partial correlation matrix, L2 regularization was applied by setting rho as 0.5 in the Ridge Regression “netmats” option. A description of the settings for the estimations is available at the following URL: http://biobank.ctsu.ox.ac.uk/crystal/refer.cgi?id=9028. To better illustrate the group-average network matrix, the nodes were clustered into 5 categories based on the full correlation matrices (Miller et al., 2016). The group-average network matrix is shown in Figure S13.

Before analysis of sex differences, we multiplied the strength of each connection by the sign of its group-mean (Smith et al., 2015). For example, where the time series data from two ICA components were positively correlated, but the valence of the connection at the level of the group was negative, the valence for that individual was determined to be negative; that is, individual valences were determined by the valence of that connection at the level of the group. In this way, the valence of the majority of participants’ connections for each node were positive, allowing us to investigate the degree to which temporal connectivity differed by sex without combining positive and negative effects and losing information on the absolute magnitude. We then tested the association of sex with the strength of connections, using the *glm* function in R. As in the other analyses, age and ethnicity were controlled by using them as covariates. Any participant without age or ethnicity information was excluded. 4,004 participants were therefore included in the analysis (mean age = 61.63, SD = 7.56; 47.65% male). To assess the importance of the nodes, we generated the weighted degree for a node by calculating the mean strength of its connections with all 54 other nodes. Full results for connection strength (partial and full correlations) and for weighted degree are provided as three separate tabs in Table S13. In that table, Cohen’s *d*-values are provided as standardised effect sizes of the sex difference in the strength of connectivity: as for the other analyses, a negative effect size means the strength of the connection was higher in males, and a positive effect size means it was higher in females.

### Cognitive testing

Cognitive testing took place at the same visit as the MRI scan. Two tests were analysed here: “fluid intelligence” (henceforth called “verbal-numerical reasoning”), and reaction time. These are described in detail in the Supplemental Materials.

### Statistical analysis

We first adjusted all variables for age and ethnicity (both of which may have been associated with differences in brain measures; Cox et al., 2016; Isamah et al., 2010; Tang et al., 2010). In some analyses, as described below, we adjusted for total brain volume and height. The adjustment techniques are described in the Supplemental Materials.

Welch’s *t*-test was used for the mean comparisons, and a variance ratio test (*F*-test) was used to assess differences in the variance between the sexes. To calculate the associated Cohen’s *d*-value for each *t*-test, we multiplied the *t*-value by 2 and divided it by the square root of the degrees of freedom. The difference between correlations for each sex was calculated using Fisher’s *r*-to-*z* transformation and a *z*-test (using the *r. test* function in the *psych* package for R; Revelle, 2016). *p*-values were adjusted, within each analysis and within each brain measure, with the False Discovery Rate correction (Benjamini & Hochberg, 1995; for example, the *p*-values for all the sex comparisons on volume were corrected separately from the *p*-values for all the sex comparisons on surface area) using the *p.adjust* function (with the “fdr” correction) for R. We used an alpha level of .05 to denote statistical significance. In an additional Bayesian analysis of the mean difference, we used the *BayesFactor* package for R (Morey & Rouder, 2015) to compute BF_10_ values from a Bayesian *t*-test (using the *ttestBF* function; see Supplemental Materials).

We used cross-sectional mediation models (in a structural equation modelling framework) to test whether the brain variables (total brain volume, grey matter volume, white matter volume, total surface area, mean cortical thickness, general fractional anisotropy, and general orientation dispersion—the latter two estimated as latent variables—each in separate models; as well as specific brain regions – see below) were significant mediators of the relation between sex and cognitive ability (either verbal-numerical reasoning score or reaction time, in separate models). We also ran multiple-mediator models that used individual brain subregions as mediators of the sex-cognitive relation, instead of overall measures. All methods for running the mediation analyses, along with the equation used to calculate a “percentage of mediation” for each brain variable, are described in the Supplemental Materials.

## Results

### Sex differences in overall and subcortical brain volumes

The subcortical structures examined were the hippocampus, the nucleus accumbens, the amygdala, the caudate nucleus, the dorsal pallidum, the putamen, and the thalamus (Figure S1). Raw volumetric sex differences are illustrated in Figure 1. The male distributions were further to the right, indicating higher means, and wider, indicating greater variance. This was confirmed by computing shift functions (Rousselet et al., 2017) for each overall and subcortical brain structure, illustrated in Figure S2. There was a substantial degree of overlap between the sexes on all measures.

**Figure 1.**
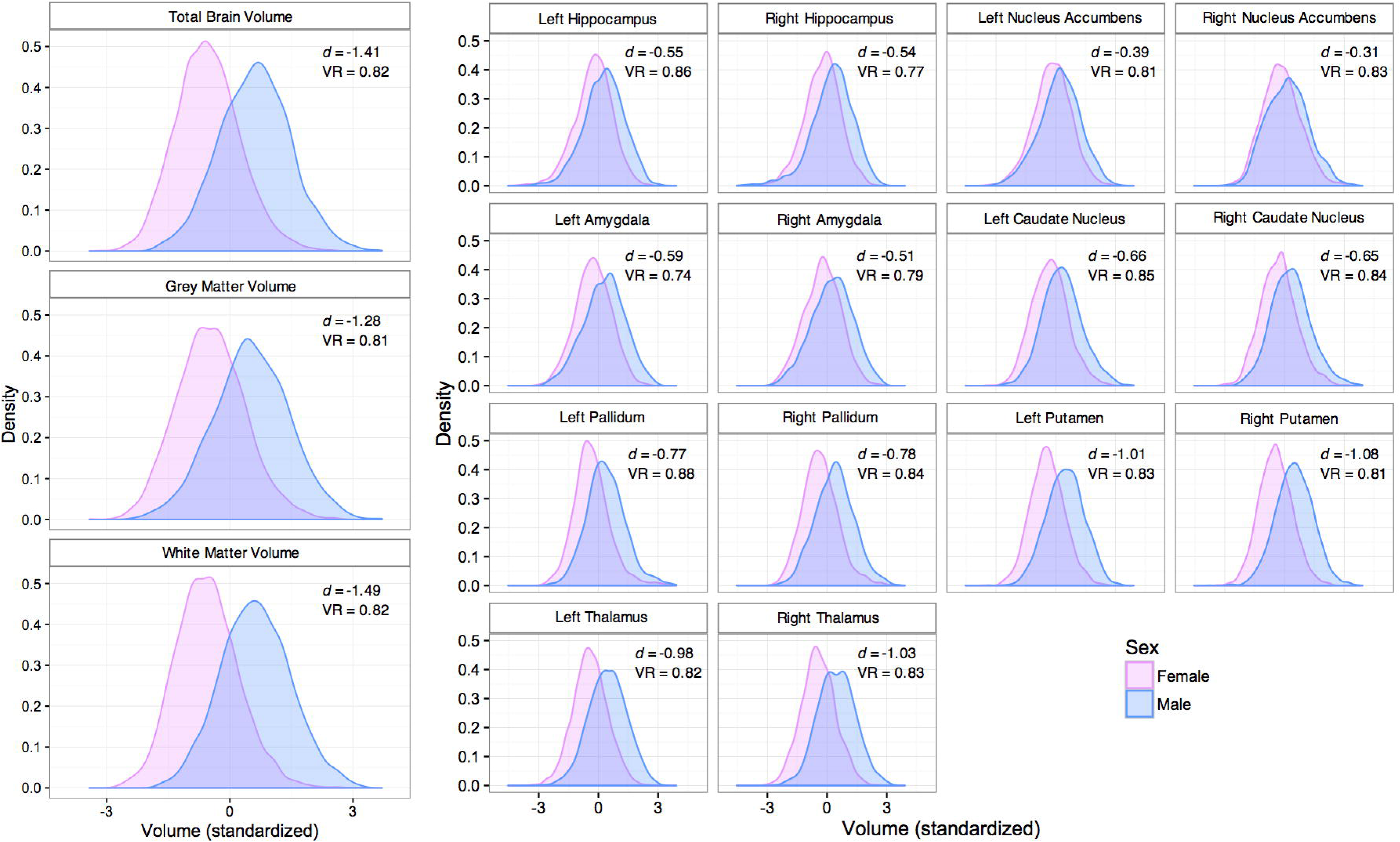
Density plots of sex differences in overall brain volumes (left section) and subcortical structures (right section). *d* = Cohen’s *d* (mean difference); VR = Variance Ratio (variance difference). All mean differences were statistically significant at *p* < 3.0×10^−25^, all variance differences were significant at *p* < .003, after correction for multiple comparisons (see Table 1).

We first tested for mean sex differences in overall cortical and subcortical brain volumes, adjusting each measure for age and ethnicity. We examined differences in total as well as grey and white matter volumes separately. Differences are shown in Table 1. We observed statistically significant sex differences (adjusted for multiple comparisons using the False Discovery Rate correction), all showing larger volume in males. In what follows, negative effect sizes indicate higher values for males, and positive effect sizes indicate higher values for females. The effect sizes ranged from small to large; for example, Cohen’s *d* = −0.39 and −0.31 for the left and right nucleus accumbens volume, respectively; −1.41, −1.28, and −1.49 for total, grey matter, and white matter volumes respectively. The average difference for the fourteen subcortical volumes was *d* = −0.70. We also tested for age-by-sex interactions, assessing whether brain measures were more strongly associated with age in males or females. This was not the case for the overall measures (adjusted *p*-values > .8). However, all of the subcortical measures except the amygdala and the caudate showed significant interactions, indicating that the age association was stronger (and the implied age trend steeper) for males. Note that the reported effect sizes come from *t*-tests on variables adjusted for age and sex, but not their interaction.

**Table 1.** Descriptive statistics with mean and variance comparisons for overall volumes, subcortical volumes, and cognitive tests.

| Measure type | Measure | Female<br>(*n* =<br>2,750) | Male<br>(*n* =<br>2,466) | Mean difference test |  |  |  | Variance Ratio test |  |
| --- | --- | --- | --- | --- | --- | --- | --- | --- | --- |
|  |  | M (SD) | M (SD) | *t* | *p* | *d* | BF<sub>10</sub> | VR | *p* |
| Overall volumes (cm<sup>3</sup>) | Total brain volume | 1115.76 (89.68) | 1233.58 (98.31) | -48.91 | ~0.00 | -1.41 | 9.57×10<sup>426</sup> | 0.82 | 6.46×10<sup>-06</sup> |
|  | Grey matter volume | 597.02 (47.78) | 643.45 (52.08) | -38.97 | 1.75×10<sup>-287</sup> | -1.28 | 1.62×10<sup>289</sup> | 0.81 | 3.60×10<sup>-06</sup> |
|  | White matter volume | 518.85 (47.89) | 589.59 (52.69) | -51.53 | ~0.00 | -1.49 | 1.47×10<sup>465</sup> | 0.82 | 7.31×10<sup>-06</sup> |
| Subcortical volumes (cm<sup>3</sup>) | Left hippocampus<sup>a</sup> | 3.73 (0.42) | 3.94 (0.46) | -18.91 | 2.69×10<sup>-76</sup> | -0.55 | 1.09×10<sup>74</sup> | 0.86 | 3.83×10<sup>-04</sup> |
|  | Right hippocampus<sup>a</sup> | 3.82 (0.42) | 4.04 (0.48) | -18.43 | 1.16×10<sup>-72</sup> | -0.54 | 7.97×10<sup>70</sup> | 0.77 | 1.16×10<sup>-09</sup> |
|  | Left accumbens<sup>a</sup> | 0.49 (0.11) | 0.53 (0.12) | -13.42 | 5.19×10<sup>-39†</sup> | -0.39 | 2.13×10<sup>36</sup> | 0.81 | 2.95×10<sup>-06</sup> |
|  | Right accumbens<sup>a</sup> | 0.40 (0.10) | 0.42 (0.11) | -10.64 | 3.82×10<sup>-26†</sup> | -0.31 | 1.04×10<sup>23</sup> | 0.83 | 4.46×10<sup>-05</sup> |
|  | Left amygdala | 1.21 (0.22) | 1.35 (0.25) | -20.04 | 5.23×10<sup>-85†</sup> | -0.59 | 4.73×10<sup>83</sup> | 0.74 | 5.89×10<sup>-12</sup> |
|  | Right amygdala | 1.18 (0.24) | 1.31 (0.27) | -17.55 | 2.16×10<sup>-66†</sup> | -0.51 | 1.60×10<sup>64</sup> | 0.79 | 1.54×10<sup>-07</sup> |
|  | Left caudate | 3.28 (0.38) | 3.54 (0.41) | -23.00 | 3.04×10<sup>-110</sup> | -0.66 | 2.70×10<sup>108</sup> | 0.85 | 2.38×10<sup>-04</sup> |
|  | Right caudate | 3.45 (0.40) | 3.72 (0.44) | -22.67 | 2.37×10<sup>-107</sup> | -0.65 | 4.08×10<sup>105</sup> | 0.84 | 4.46×10<sup>-05</sup> |
|  | Left pallidum<sup>a</sup> | 1.69 (0.21) | 1.85 (0.22) | -26.64 | 4.87×10<sup>-145†</sup> | -0.77 | 2.19×10<sup>143</sup> | 0.88 | .002 |
|  | Right pallidum<sup>a</sup> | 1.74 (0.20) | 1.89 (0.22) | -26.96 | 3.82×10<sup>-148†</sup> | -0.78 | 8.59×10<sup>146</sup> | 0.84 | 1.03×10<sup>-04</sup> |
|  | Left putamen<sup>a</sup> | 4.61 (0.50) | 5.07 (0.56) | -34.72 | 1.73×10<sup>-234†</sup> | -1.01 | 1.29×10<sup>235</sup> | 0.83 | 1.46×10<sup>-05</sup> |
|  | Right putamen<sup>a</sup> | 4.64 (0.49) | 5.13 (0.55) | -37.13 | 4.76×10<sup>-264†</sup> | -1.08 | 3.02×10<sup>265</sup> | 0.81 | 1.98×10<sup>-06</sup> |
|  | Left thalamus<sup>a</sup> | 7.54 (0.64) | 8.11 (0.72) | -33.73 | 7.76×10<sup>-223</sup> | -0.98 | 1.50×10<sup>223</sup> | 0.82 | 1.34×10<sup>-05</sup> |
|  | Right thalamus<sup>a</sup> | 7.34 (0.62) | 7.92 (0.69) | -35.76 | 2.42×10<sup>-247</sup> | -1.03 | 6.62×10<sup>247</sup> | 0.83 | 4.46×10<sup>-05</sup> |
| Cognitive tests | Verbal-numerical reasoning (max. score 13) | 6.80 (2.10) | 7.14 (2.13) | -6.21 | 5.77×10<sup>-10</sup> | -0.18 | 6.94×10<sup>6</sup> | 0.97 | .451 |
|  | Reaction time (ms) | 590.37 (98.04) | 574.71 (100.71) | -7.63 | 2.71×10<sup>-14</sup> | -0.21 | 1.30×10<sup>11</sup> | 0.92 | .033 |

*Note:* Means and SDs are shown prior to adjustment for age and ethnicity; statistical tests are performed after this adjustment. Reaction Time is shown here in raw millisecond units, but was reverse-scored for analysis so that higher scores indicated better performance. Negative *t*- and *d*-values mean higher male means. VR = Variance ratio (values < 1 indicate greater male variance). *p*-values for brain variables corrected for multiple comparisons using the False Discovery Rate correction. BF<sub>10</sub> = Bayes Factor indicating the probability of the alternative hypothesis (that there is a sex difference) compared to the null hypothesis (that there is no sex difference). <sup>a</sup> = significant age-by-sex interaction. <sup>†</sup> = sex difference in subcortical region still significant after adjustment for total brain volume (see Table S1).

We tested whether sex differences in the subcortical measures were accounted for by the substantial difference in total brain volume. We regressed each subcortical variable on total brain volume, testing the residuals for sex differences. After this adjustment, there were no longer statistically significant differences in the hippocampus, caudate nucleus, or thalamus (all *p*_adj_-values > 0.60, absolute *d*-values < 0.03; Table S1). There remained differences in each of the other measures, with attenuated effect sizes (average *d* for significant differences after adjustment = 0.17). Females had greater nucleus accumbens volume after adjustment for total brain volume (*d* = .08, *p*_adj_ = .07 for left accumbens; *d* = 0.10, *p*_adj_ = .003 for right). Overall, the majority of the sex differences in specific subcortical structures appeared to be linked to the total brain size (average attenuation of *d*-values for subcortical structures = 85.0%). We also ran analyses adjusting for height, since overall body size may have influenced these differences (as expected, males were substantially taller: *d* = −2.15). This attenuated all of the *d*-values (average attenuation across global and subcortical measures = 71.3%), but males still showed significantly larger volumes for all regions except the nucleus accumbens (Table S1). For example, post-adjustment *d*-values were −0.42 for total brain volume, −0.31 for grey matter volume, and −0.47 for white matter volume.

As shown in Table 1, there were statistically significant variance differences in all overall cortical and subcortical brain volumes, with males showing greater variance; the average variance ratio for overall volumes and subcortical volumes was 0.82 (variance ratios <1.00 indicate greater male variance). After adjusting for total brain volume or height, the variance differences reported in Table 1 remained relatively unchanged (see Table S1).

### Sex differences in subregional brain volume, surface area, and cortical thickness

Using Freesurfer to parcellate cortical regions according to the Desikan-Killiany neuroanatomical atlas (Desikan et al., 2006; Figure S3), we tested for sex differences in volume, surface area, and cortical thickness across 68 cortical subregions. As with the analyses above, we adjusted all subregions for age and ethnicity; *p*-values were also adjusted within each measure type using the False Discovery Rate correction. The results are illustrated in Figure 2A (see also Table S2 for means, standard deviations, and difference tests for volume, surface area, and cortical thickness across all cortical regions).

**Figure 2.**
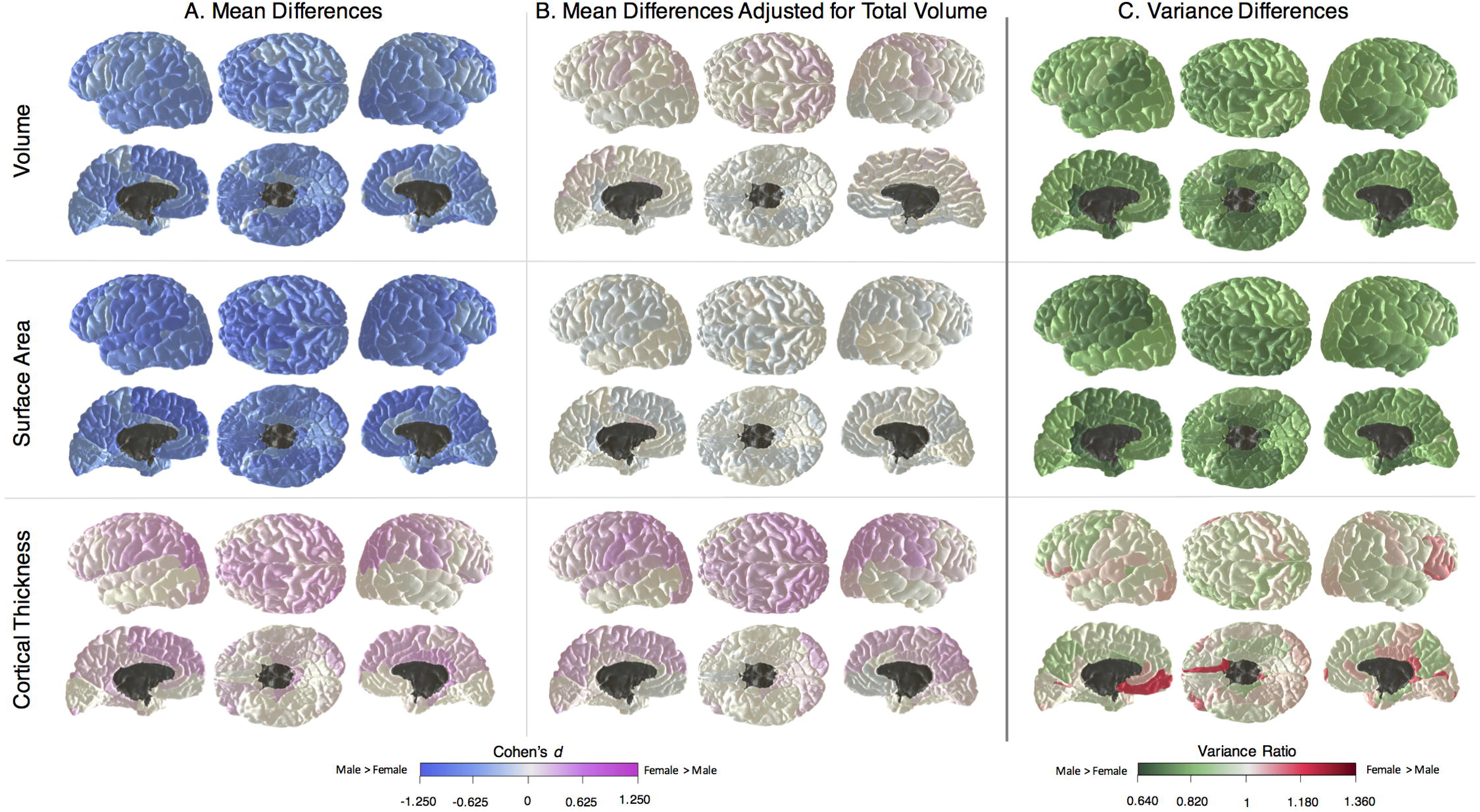
Sex differences across the subregions in volume, surface area, and cortical thickness. Shown are A) mean differences, B) mean differences adjusted for total brain volume, and C) variance differences. Variance differences corrected for total brain volume were near-identical to those shown in C); see Figure S5. See Figure S3 for subregional atlas.

Males showed larger brain volume across all cortical subregions. The sex differences were statistically significant in every subregion, ranging in size from small (*d* = −0.24 in the right temporal pole) to large (*d* = −1.03 in the right insula). The mean *d*-value across all subregions was −0.67 (*p*_adj_-values < 9.00×10^−13^). Even larger differences, all favouring males, were observed for surface area; these ranged from moderate (*d* = −0.43 in the left caudal anterior cingulate) to large (*d* = −1.20 in the left superior frontal region). The mean *d*-value across all subregions was −0.83 (all *p*_adj_-values < 2.00×10^−36^).

Cortical thickness displayed a different pattern. Unlike volume and surface area, females had thicker cortex across almost the entire brain. The only area where males showed a statistically significantly thicker cortex was the right (but not left) insula, and the difference was small (*d* = 0.14). In all other areas, there was either no significant thickness difference (20/68 areas), or a statistically significant difference favouring females. The mean *d*-value in the 47 areas that reached statistical significance after multiple-comparisons correction was 0.22, ranging from *d* = 0.07 in the right rostral middle frontal region to *d* = 0.45 in the left inferior parietal region. Higher female cortical thickness was generally not found in the temporal lobe (except the parahippocampal gyrus) or in the medial orbitofrontal regions. In some regions there appeared to be converse differences: in the motor and somatosensory regions in the parietal lobe, the frontal pole, and the parahippocampal gyrus, females showed relatively higher thickness but males showed relatively higher volume and surface area. In the superior temporal lobe and orbitofrontal regions, males showed relatively higher volume and surface area, but there was no particular sex difference in thickness.

We also tested age-by-sex interactions for each of the three variables (Table S2). After multiple-comparisons correction, only two interactions were significant: the left and right superior frontal regions showed significantly stronger relations with age in males. That is, males may have had steeper volume decline in this region bilaterally with age. There were no statistically significant age-by-sex interactions for surface area or cortical thickness.

We next adjusted the subregional cortical measures for total brain volume. As shown in Figure 2B (and Table S3), 11 regions were still significantly larger in volume for males. However, the majority of regions (44/68) no longer showed significant volume differences (−0.08<*d*<0.08, mean *d* = −0.01, all *p*_adj_-values > .34). There were also 13 regions where females now had a significantly larger volume relative to the total size of the brain, the largest being the right superior parietal (*d* = 0.21). For surface area, males were larger in 36/68 areas after TBV adjustment (the largest being in the left isthmus cingulate; *d* = −0.22), and females were larger in one (the left caudal anterior cingulate, *d* = 0.11). For cortical thickness, after correction for total brain volume there were still significant differences favouring females in 46/68 regions (0.07<*d*<0.41, mean *d* = 0.21, all *p*_adj_-values < .03), and four regions with differences favouring males (−0.14<*d*<-0.09, mean *d* = −0.12, all *p*_adj_-values < .02). Next, we adjusted the cortical subregional measures for height (Table S4). For volume, all of the comparisons were still significant, but with reduced effect sizes (−0.33<*d*<-0.07, mean *d* = −0.19, all *p*_adj_-values < .05); this was the same for surface area (−0.35<*d*<-0.10, mean *d* = −0.25 all *p*_adj_-values < .002). For thickness, there were 34/68 regions that were still significantly thicker in females (0.06<*d*<0.20, mean *d* = 0.12, *p*_adj_-values < .05), and one favouring males (the left entorhinal cortex, *d* = −0.08).

Variance differences across the three structural measures are illustrated in Figure 2C. For volume and surface area, males showed significantly greater variance than females across almost all brain regions. The volume variance ratio was significant in 64/68 regions, ranging from 0.88 in the right temporal pole to 0.67 in the left isthmus cingulate, with all *p*_adj_-values < .031 after correction. The surface area variance ratio was significant in 66/68 regions, ranging from 0.88 in the left pars orbitalis to 0.65 in the left isthmus cingulate, all *p*_adj_ - values < .018 after correction. For cortical thickness (Figure 2C), there were no significant variance differences in any region (all *p*_adj_-values > .14) except one: females showed significantly greater variance in the thickness of the left medial orbitofrontal cortex (VR = 1.19, *p*_adj_ = .01). As can be observed from Figure S5 (and Table S3), controlling for total brain volume made only a negligible difference to the pattern of variance ratios reported above.

We tested whether the regions showing larger mean differences were also those with larger variance differences, by correlating the vector of *d*-values with the vector of VRs for each brain measure. As shown in Figures S6 and S7, there was no clear correspondence between mean and variance: in the unadjusted analysis, mean and variance were correlated at *r* = .51 for volume, but there were smaller correlations for surface area and thickness (*r*-values = .25 and −.06, respectively). Adjusted for TBV, all three correlations were relatively weak (*r*-values = .22, .03, and −.25 for the three brain measures respectively).

To verify whether the pattern of results across the cortical mantle was agnostic to the gyral boundaries of the Desikan-Killiany atlas, we conducted a supplemental analysis, testing sex differences using a vertex-wise approach, the results of which are shown in Figures S8 (for mean differences) and S9 (for variance differences). This precisely replicated the subregional atlas-based results.

### Sex differences in white matter microstructure

We tested sex differences in 22 white matter tracts. We focused on two white matter microstructural properties that had previously been shown to demonstrate differences between males and females in the initial release of UK Biobank imaging data (Cox et al., 2016). The first was fractional anisotropy (FA), an index of the directionality of water diffusion through the white matter. The second was orientation dispersion (OD), a NODDI measure of white matter tract complexity. For FA, there were generally higher values in males, particularly in the cortico-spinal tract (*d* = −0.54) and the acoustic radiation (*d* = −0.51). The average difference across tracts was *d* = −0.19. OD was higher in all tracts for females (average *d* = 0.30). These mean differences are shown in Figure 3, and fully reported in Tables S5 and S6.

**Figure 3.**
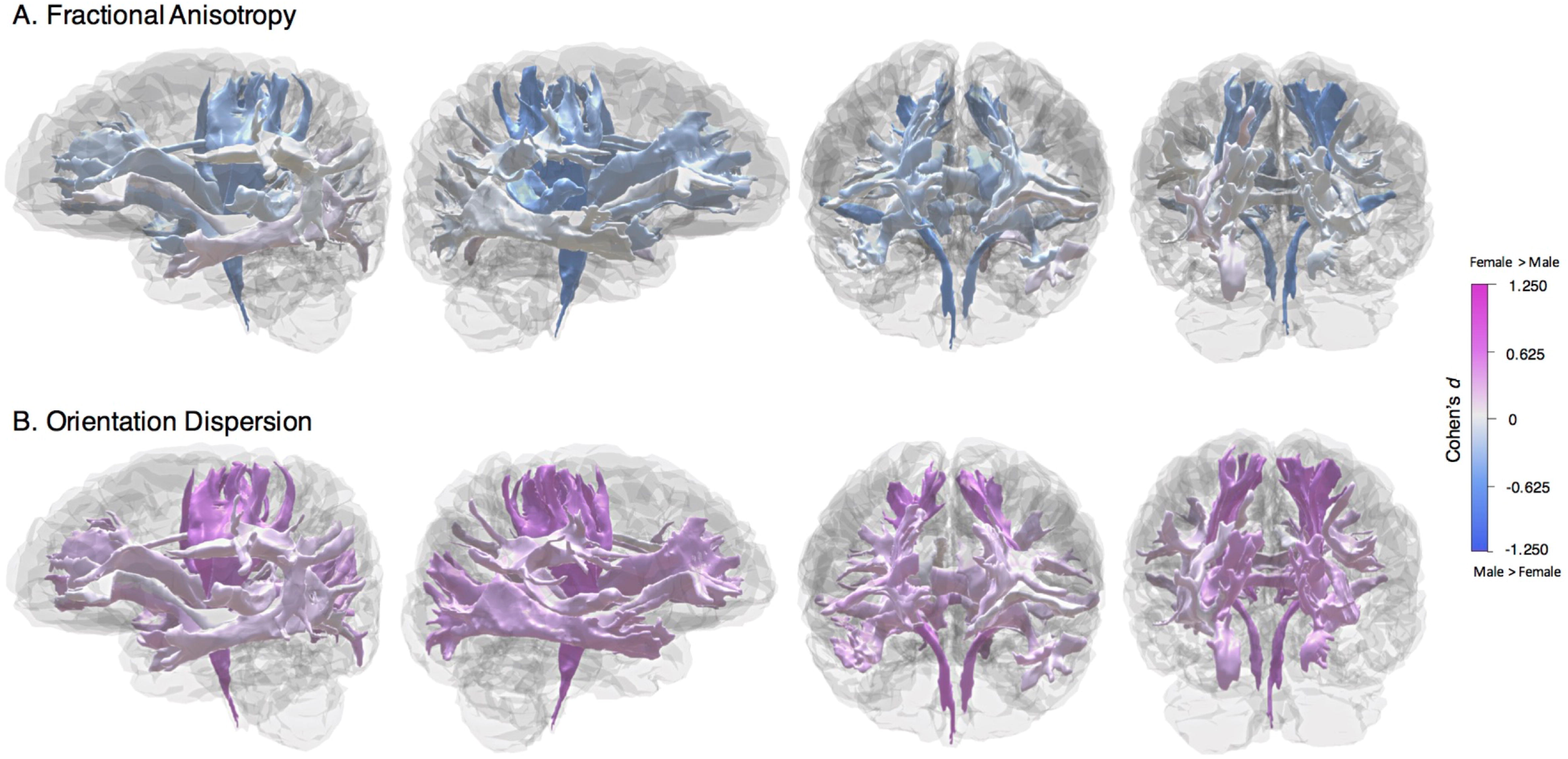
Mean sex differences in white matter microstructural measures A) fractional anisotropy and B) orientation dispersion across 22 white matter tracts. For both measures, numerically the largest effect was found in the right cortico-spinal tract. See Figure S4 for tract atlas.

Variance differences are illustrated in Figure S10 (see also Tables S5 and S6). Generally, there was greater male variance in FA (average VR = 0.92); however, there was substantially greater female variance in the cortico-spinal tract in particular (VR = 1.17, *p* = .0003). For OD, the only tract that showed a significant variance difference following FDR correction was the left superior thalamic radiation, where males showed greater variance (VR = 0.79).

### Relation of neurostructural differences to cognitive differences

We linked the structural brain differences to scores on two cognitive tests taken at the time of the imaging visit: verbal-numerical reasoning and reaction time (see Method). Descriptive statistics for the cognitive tests are shown in Table 1. Note that we coded (reflected) both tests so that higher scores indicated better performance. The test scores correlated positively, but weakly (*r* = .12). Males had a slightly higher mean score than females on verbal-numerical reasoning (*d* = −0.18) and slightly faster mean reaction time (*d* = −0.22); there was no significant variance difference for verbal-numerical reasoning (VR = 0.97, *p* = .45), though males had marginally more variance in reaction time (VR = 0.92, *p* = .03).

As a first step toward the mediation analyses, we correlated performance on the two cognitive tests with the overall brain measures in the full sample (Table S7), and in two randomly-selected sample halves separately (Table S8). The sample was split in this way to avoid overfitting and assess the replicability of the results. We then ran the same correlations across all the brain subregions, for volume, surface area, and cortical thickness (Table S9). These correlations were generally small, with all brain-cognitive *r*-values <.20. We compared the size of the correlations across the sexes; after multiple comparisons correction, there were no significant sex differences in these correlations. Thus, there was no evidence in the present analysis for sex differences in how regional brain structure related to the two measured cognitive skills.

Next, we tested the extent to which the mean cognitive differences were mediated by any of the overall brain measures (total, grey, and white matter volumes, total surface area, mean cortical thickness, or general factors of FA or OD). We ran a separate model, illustrated in Figure S11, for each brain measure. Results are displayed in Tables S10 and S11 for verbal-numerical reasoning and reaction time, respectively. For verbal-numerical reasoning, the sex difference in test scores was mediated substantially by brain volume measures and by surface area (all mediation percentages >82%). Cortical thickness showed far smaller mediation percentages (7.1% and 5.4% in the two sample halves, respectively). For reaction time, total brain and white matter volumes had mediation percentages >27%, but the other measures all produced smaller percentages (<15.3%), particularly mean cortical thickness (mediating <3% of the variance).

Finally, we tested which brain subregions were most important in explaining the mediation of the sex-cognitive relation, by running mediation models that included multiple individual regions as mediators. These variables were selected for their association with the cognitive ability in question (again, either verbal-numerical reasoning or reaction time) using LASSO regression models (see Method for details). The percentage of mediation for each selected region is illustrated in Figure 5 (see Table S12 for full results). For verbal-numerical reasoning, the volume and surface area of the superior temporal region mediated the largest amounts of variance (29.1% and 18.4% in their respective models), with other relatively substantial contributions coming from the precuneus and insula for volume, and the pars opercularis and rostral middle frontal regions for surface area. For the cortical thickness predictors, and for the outcome of reaction time, as expected on the basis of the overall mediation results reported above, few of the regions showed substantial mediation (there was some mediation by the volume of frontal regions; at most 7.3% by the frontal pole).

### Sex differences in resting-state functional connectivity

For our final set of analyses, we examined sex differences in resting-state functional MRI (rsfMRI) responses within a number of functional networks. The connections between each pair of functional networks were estimated and then transformed into measures of strength (see Method). We found that 54.7% (811 of 1,485) of network connections showed a statistically significant sex difference (absolute *β*-values= 0.071-0.447 for females; 0.071-0.519 for males). A map showing the strengths of the connections between the 55 network nodes, and whether the difference was stronger in males (blue) or females (red) is provided in Figure 5A (see also Table S13). The strength of connectivity between sensorimotor, visual, and rostral lateral prefrontal areas was absolutely higher in males than females (see the cluster of brain regions with orange numerals in Figure 5A), whereas the strength of connectivity within the default mode network (DMN; cluster of regions with red numerals in Figure 5A) was absolutely higher in females than males.

**Figure 4.**
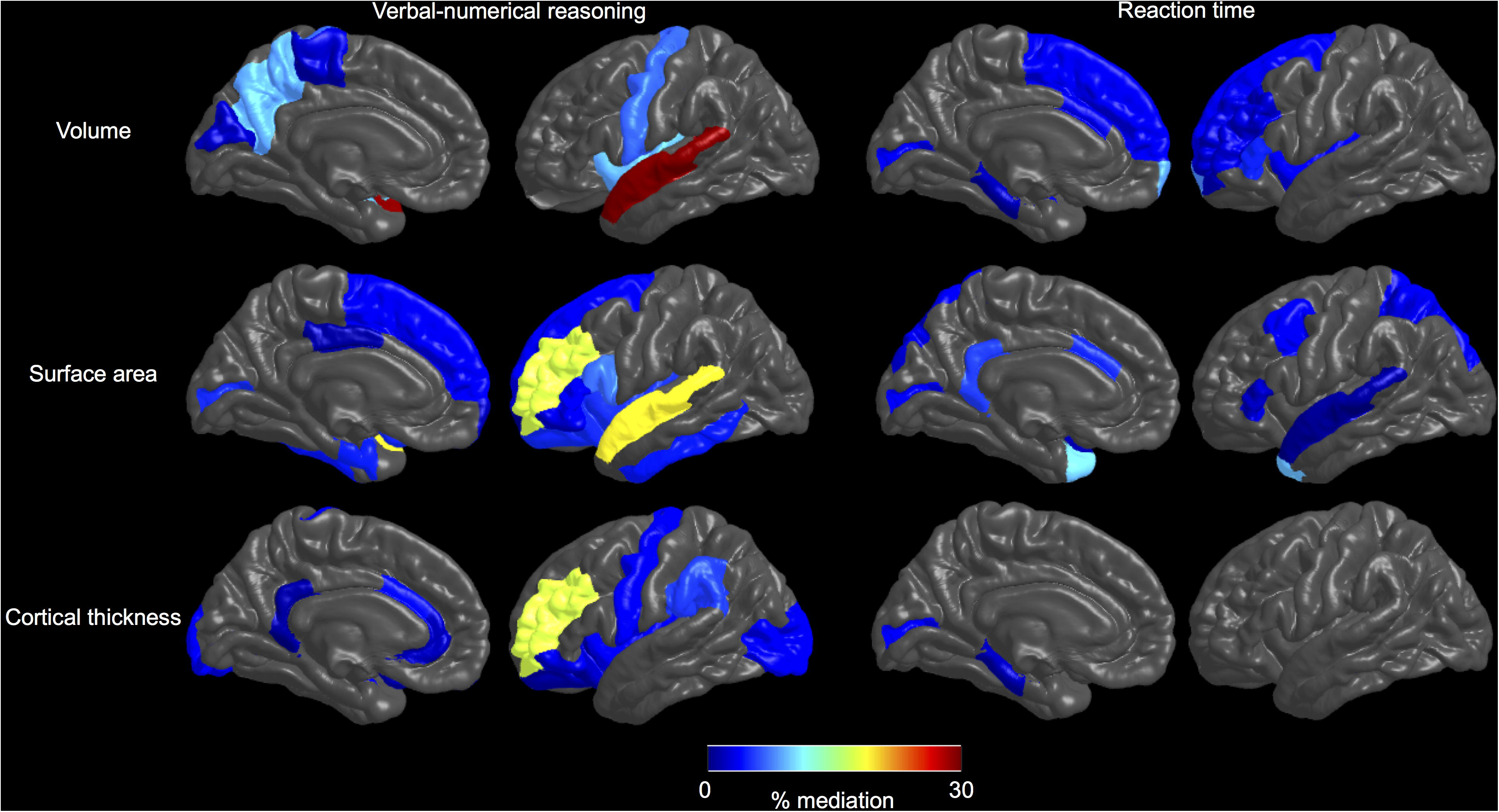
Percentage of the sex-cognitive relation mediated by each of the brain regions selected in a LASSO model to be linked to either verbal-numerical reasoning (left column) or reaction time (right column). Results for volume, surface area, and cortical thickness are shown in each row. Regions were averaged across the hemispheres; thus only a medial and lateral view for each measure and each cognitive test is shown.

**Figure 5.**
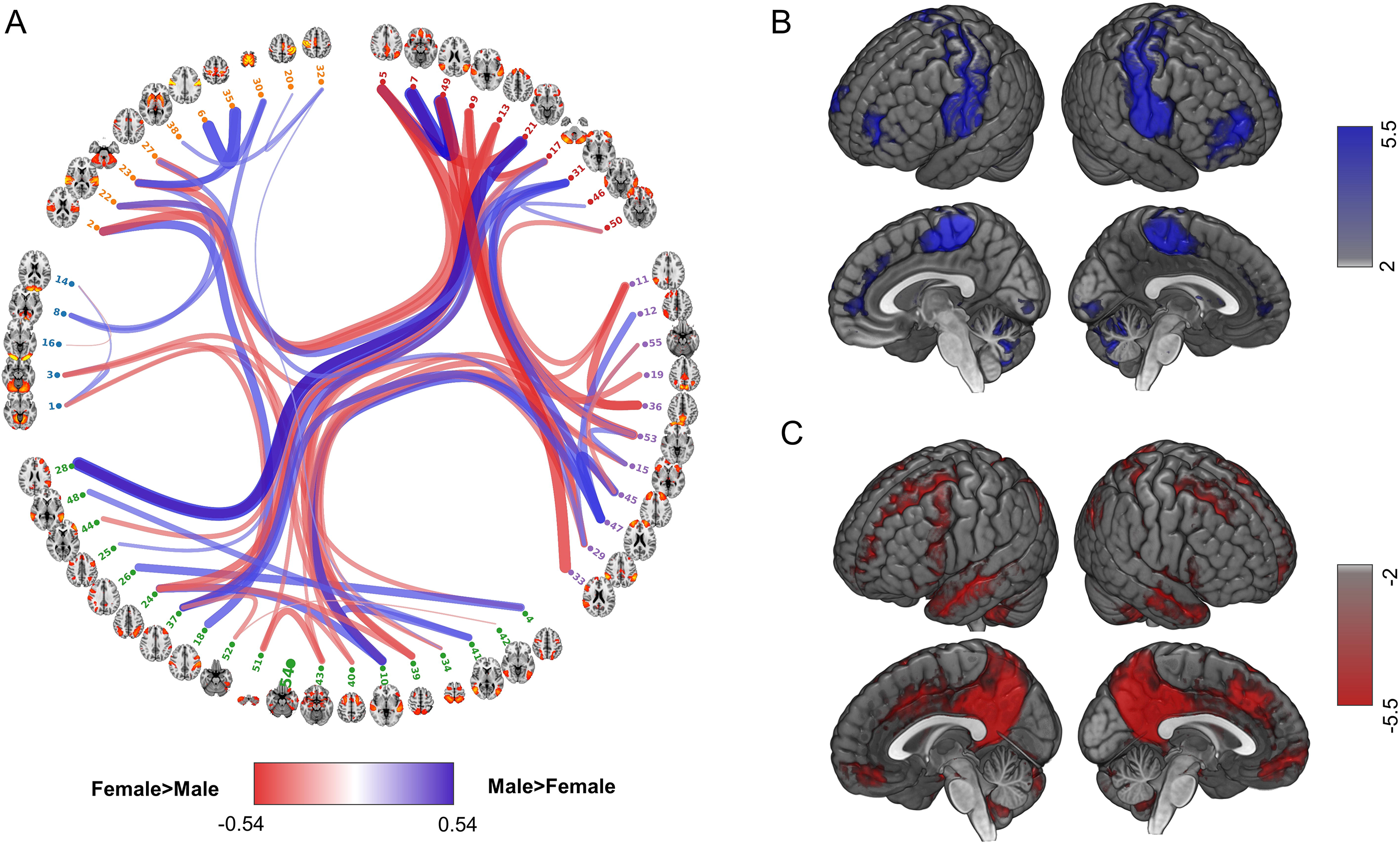
Results for resting-state *ƒ*MRI connectivity and weighted degree of nodes. A) Spatial maps for individual connections. Colours and line thickness represent the effect sizes of sex on the strength of connections (red = stronger in females; blue = stronger in males; darker/thicker = larger effect size). Only effect sizes (Cohen’s *d*) larger than ±0.2 are shown. Nodes were clustered into five categories using FSLnets based on their group-mean full-correlation matrix (yellow/orange: sensorimotor network; red: default mode network; purple: salience network and executive control network; green: dorsal attention network; blue: visual network). B) and C) Weighted degrees of nodes with higher values in males and females, respectively. The spatial maps of significant group-ICA nodes were multiplied by the effect size of the sex correlation. In order to show the regions with the largest associations with sex, only regions that had intensity over 50% of the whole-brain peak value are presented. See Table S14 for values for each connection and for each node’s weighed degree.

To further analyse these functional sex differences, we calculated the mean strength of all 54 connections to each individual node, producing a “weighted degree” statistic. Sex differences in weighted degree are shown in Figures 5B and 5C. Males showed stronger weighted degree than females in bilateral sensorimotor areas, the visual cortex, and the rostral lateral prefrontal cortex. Females showed stronger weighted degree than males in cortical areas comprising the DMN: the bilateral posterior cingulate cortex/precuneus, the dorsal anterior cingulate cortex, medial prefrontal cortex, temporo-parietal junction, anterior temporal lobe, medial temporal lobe (e.g. hippocampus and surrounding areas), and some cerebellar regions (see Tables S13 and S14).

## Discussion

In a single-scanner sample of over 5,000 participants from UK Biobank, we mapped sex differences in brain volume, surface area, cortical thickness, diffusion parameters, and functional connectivity. One main theme of the neurostructural results was that associations with sex were global. Males generally had larger volumes and surface areas, whereas females had thicker cortices. The differences were substantial: in some cases, such as total brain volume, more than a standard deviation. We also found that volume and surface area mediated nearly all of the small sex difference in reasoning ability, but far less of the difference in reaction time. For white matter microstructure, females showed lower directionality (FA) and higher tract complexity (OD); white matter microstructure was a poor mediator of the cognitive sex difference. Resting-state fMRI analyses also revealed a global effect: around 54% of connections showed a sex difference. These differences clustered around specific networks, with stronger connectivity in females in the default mode network and stronger connectivity in males between unimodal sensory and motor cortices as well as high-level cortical areas in the rostral lateral prefrontal cortex. Overall, for every brain measure that showed even large sex differences, there was always overlap between males and females (see Figure 1).

The principal strengths of the present study are its sample size (providing sensitivity for the identification of small effects with high statistical power), the wide range of MRI modalities, and the consideration of both mean and variance differences. Given the surfeit of small-*n* studies in neuroscience (Button et al., 2013; Nord et al., 2017), it is of great importance to test hypotheses in large, well-powered samples, especially given that many neural sex differences are modest in size (Joel et al., 2015). Here, we had excellent statistical power to find small effects in brain subregions, providing a robust and detailed analysis. For our subregional analysis, we had a far larger sample size than the most recent meta-analysis (Ruigrok et al., 2014). In contrast to that meta-analysis—which found greater volume for females in areas such as the thalamus, the anterior cingulate gyrus, and the lateral occipital cortex—our study found no brain subregions where females had a larger volume than males. The reason for this may be the more restricted age range of the participants in our study (sex may relate differently to the brain at different ages, as we found for several brain regions in our age-by-sex interaction analyses, and as was found in a previous developmental study of children and adolescents; Gennatas et al., 2017) or, more likely, study size and heterogeneity: the data for section of the meta-analysis on regional volumes came from many separate studies, on separate scanners, generally with small sample sizes (many with *n* < 100), whereas our contrasts were based on one very large, single-scanner study.

The higher male volume in our study appeared largest in some regions involved in emotion and decision-making, such as the bilateral orbitofrontal cortex, the bilateral insula, and the left isthmus of the cingulate gyrus (Craig, 2009; MacPherson et al., 2015; Ochsner & Gross, 2005; Wager et al., 2008; note that the insula showing the largest sex difference is consistent with a recent large-scale study of children and adolescents (Gennatas et al., 2017) – it appears this region retains its substantial sex difference into later life), but also areas such as the right fusiform gyrus. For surface area, which showed an even larger difference favouring males, the regions that showed the largest effects were broadly areas involved in the hypothesized intelligence-related circuit in the “P-FIT” model (Jung & Haier, 2007): for example, the bilateral superior frontal gyri, the bilateral precentral gyri, the left supramarginal gyrus, and the bilateral rostral middle frontal areas. However, some of the regions involved in this theorized circuit were also larger, in terms of thickness, for females. For instance, the bilateral inferior parietal regions were the regions with numerically the largest difference favouring females in cortical thickness. Our finding that the cortex was thicker for females is consistent with a number of previous, smaller studies (e.g. Luders et al., 2006; Lv et al, 2010; Sowell et al., 2007; van Velsen et al., 2013), though our greater statistical power allowed us to find smaller differences in thickness across the cortex.

Whereas previous work has found some white matter regions where fractional anisotropy was higher for females (Kanaan et al., 2012; Dunst et al., 2014), we found that males showed higher FA in 18 of the 22 tracts we examined. FA also generally showed greater variance in males. On the other hand, higher orientation dispersion was found for females in all tracts. Unexpectedly, higher OD was found to be related to lower cognitive performance on the two tests examined here. Since OD is a relatively new measure of white matter microstructure (Daducci et al., 2015), further work should aim to clarify its behavioural correlates. The fact that (as described in the Method section) measurement invariance did not hold across the sexes for the latent variables of FA and OD, indicating that the tract-specific measurements may be assessing somewhat different latent variables in each sex, may also be of interest for future researchers examining general-level indicators of white matter microstructure.

The issue of adjusting for overall brain size in analyses of sex differences (Rippon et al., 2014) was addressed in each of our macrostructural analyses. As can be seen comparing Figures 2 and 3, after this adjustment, the higher male volume and surface area was substantially reduced, often to non-significance. For those latter brain regions, this implies that the sex difference was general and that the larger volume or surface area was a byproduct of the overall larger male brain. However, for some regions, especially for surface area (particularly in areas such as the left isthmus of the cingulate gyrus and the right precentral gyrus), males still showed a significantly higher measurement, indicating specific sex differences in the proportional configuration of the cortex, holding brain size equal. Most interestingly, for some areas (for example the right insula, the right fusiform gyrus, and the left isthmus of the cingulate gyrus), the difference was reversed after adjustment, with females showing significantly larger brain volume.

A recent meta-analysis of sex differences in amygdala volume (Marwha et al., 2016) found that, although males showed larger raw volume, after correction for total brain volume there was no longer an appreciable sex difference. However, in our study the amygdala was significantly, but modestly, larger in males even after adjusting for total brain volume (*d* = 0.18 bilaterally). The heterogeneity in the methods of the studies being meta-analysed may have led to the divergent conclusion from our single-sample study. With regard to the hippocampus, however, we found results consistent with another recent meta-analysis (Tan et al., 2016): there were no longer significant sex differences after adjustment for total brain volume (this was also the case for the thalamus and caudate). We recommend that future studies perform comparisons both before and after adjusting for total volume, since these results pertain to quite different questions.

One question that could not be addressed using the current data regards the underlying bio-social causes, ultimate or proximate, for the sex differences that we observed. Many variables were collected in UK Biobank that might be linked to the sex differences observed here (and may be proximal causes of them) but our intention in the present study was to characterise, not necessarily explain, these differences: future research should investigate more targeted hypotheses of the causes of the differences. Sex differences in brain structure are observed early in the life course (e.g. Knickmeyer et al., 2014), though this does not imply that the pattern of adult differences we observed is necessarily the same as is found in childhood. The literature on developmental sex differentiation of the brain highlights influences of factors such as sex hormones that were not analysed in the present study (Lombardo et al., 2012; McCarthy & Arnold, 2011; McEwen & Milner, 2017). Likewise, understanding the potential neurobiological effects of social influences during development (Dawson et al., 2000) was beyond the scope of our research and our dataset.

Our analysis also focused on sex differences in variability. Here, for the first time in an adult sample, we directly tested sex differences in the variance of several brain measures, finding greater male variance across almost the entire brain for volume, surface area, and white matter fractional anisotropy, but only patchy and inconsistent variance differences for cortical thickness and white matter orientation dispersion. One potential candidate to explain greater male variability across multiple phenotypes is the hypothesized ‘female-protective’ mechanism involving effects of the *X* chromosome (Craig et al., 2009; Johnson et al., 2009; Reinhold & Engqvist, 2013), or other protective factors that might “buffer” females from potential deleterious consequences of rare genetic mutations (Jacquemont et al., 2014; Robinson et al., 2013). Such explanations are speculative at present; as studies like UK Biobank release even larger amounts of data on individuals who have both neurostructural and genotype data, researchers will be able to perform well-powered tests of these hypotheses.

Using the (limited) data on cognitive abilities available in our sample, we tested whether the data were consistent with any consequences of brain structural differences in terms of ability differences. There were only weak correlations between brain variables and the cognitive tests, and these associations did not differ by sex (consistent with a prior meta-analysis on the link between brain volume and intelligence; Pietschnig et al., 2015). Mediation modelling suggested that, for verbal-numerical reasoning, a very large portion (up to 99%) of the modest sex difference was mediated by brain volumetric and surface area measures. Smaller fractions (up to 38%) of the modest link between sex and reaction time could be explained by volume or surface area. Perhaps unexpectedly, given evidence and theory linking white matter microstructure to cognitive processing speed (Bennett & Madden, 2014; Penke et al., 2012), white matter microstructural measures only mediated a small proportion of the sex difference in reaction time (this may have been due to weaknesses in this cognitive measure; see below). Cortical thickness had trivial mediating effects compared to volume and surface area: no more than 7.1% of the sex-cognitive relation was mediated by thickness in any analysis. With our multiple-mediator models, we built a map of which brain regions were most relevant in this mediation of the sex-cognitive reaction (Figure 4). Overall, the data were consistent with higher volume and cortical surface area—but not cortical thickness or microstructural characteristics—chiefly in the superior temporal region, but also spread across multiple other regions to a lesser extent, being of particular relevance to sex differences in reasoning (but not reaction time).

Sex differences in intrinsic functional connectome organization also revealed results that corroborate and extend prior work. We successfully replicated the results from the 1,000 Functional Connectomes dataset (an entirely separate dataset) – that is, we found female>male connectivity within the default mode network and some evidence for male>female connectivity in sensorimotor and visual cortices (Biswal et al., 2010). The higher female connectivity within circuits like the DMN may be particularly important, given that DMN regions are typically considered as the core part of the “social brain” (Kennedy & Adolphs, 2012). Whether such an effect can help explain higher average female ability in domains like social cognition (Gur et al., 2012), and whether such functional differences can be integrated with differences in the structural connectome (Ingalhalikar et al., 2014), remains to be seen. Finally, recent work has shown that intrinsic functional connectome organization can be parsimoniously described as a small number of connectivity gradients (Margulies et al., 2016). The most prominent connectivity gradient has at one pole the DMN and at the other unimodal sensory and motor cortices. The observed pattern of sex differences in functional connectome organization observed here appears to recapitulate the two main poles of that principal connectivity gradient (Margulies et al., 2016). One potential way of describing the biological significance of these functional sex differences is that mechanisms involved in shaping sex differences (biological, cultural, or developmental) may influence this principal connectivity gradient; the result, which should be explored in future investigations of brain sex differences, may be the multiple network differences found in the present study.

### Limitations

The UK Biobank sample was selective. It covered only one part of the life course (from approximately 45 to 75 years of age). Many of the female participants might have been undergoing, or have undergone, menopause; this (or associated Hormone Replacement Therapy) might exert modest effects on the structure of some regions of the brain (Zhang et al., 2016), effects which may themselves change with increasing age. In addition, UK Biobank had a very low response rate to invitations to participate (5.47% in the full sample of ~500,000; Allen et al., 2012). We would thus expect the individuals studied here would not be fully representative of males and females from the general UK population. This was the case for education: individuals with college or university degrees were over-represented (see Method), though the male:female education ratio itself appeared representative. Although we adjusted for the effects of age, it should also be noted—as for any study with a wide age range—that there was substantial variation in the birth date of the participants, undoubtedly leading to different (unmeasured) social experiences during their development. On the topic of age adjustment, it should also be noted that we adjusted for linear effects of age, whereas some variables may have nonlinear trends (although we would not expect this to affect the sex difference in these variables to a substantial extent). A final issue of representativeness concerns clinical outcomes. Although we noted above that there is much interest in sex-differential patterns of psychiatric disorder diagnoses, the unrepresentativeness of UK Biobank extends to generally low rates of such disorders in general in the sample. For this reason, we did not attempt to link the MRI sex differences observed here to clinical diagnoses, though studies of normal-range variation in traits linked to psychiatric disease (such as neuroticism, a known risk factor for Major Depressive Disorder; Kotov et al., 2010), may produce more fruitful results.

Caution should be taken in interpreting the results of the analyses involving the cognitive tests (the mediation analyses in addition to the correlations). Whereas previous, representative studies (e.g. Johnson et al., 2008) have found no mean difference, but a variance difference, in general cognitive test performance, the tests examined here showed mean differences but no strong variance differences. This may be due to problems of sample representativeness (Dykiert et al., 2009), or due to the tests tapping specific cognitive skills rather than general ability (Burgleta et al., 2012). The cognitive measures were relatively psychometrically poor compared to a full IQ assessment: the verbal-numerical reasoning had only 13 items, and the reaction time test had only 4 trials that counted towards the final score (see Lyall et al., 2016, for analyses of the reliability of these tests). Although the tests— particularly verbal-numerical reasoning—have some external validity (Hagenaars et al., 2016), the above issues mean that the cognitive analyses reported here should be considered preliminary. Fuller cognitive testing, currently underway in UK Biobank, will allow a more comprehensive exploration. Studies that use tests where males or females are known to show higher average scores (such as 3D mental rotation tests, which generally show higher scores in males; Maeda & Yoon, 2013), would potentially allow for more informative results. In addition, cross-sectional mediation models of observational data, such as those used here, are inherently limited: they cannot address causal relations between variables. The models were simple, including only three main variables (sex, the brain measure, and cognitive ability; Figure S11). More complex models, using longitudinal data and latent variables derived from multiple cognitive tests, should be specified in future research.

Finally, although this study used a wide variety of neuroimaging measures, it should be noted that these were but a small selection of the possible modalities that we could have investigated, and that studies should address in future. Other diffusion and NODDI measures of white matter microstructure such as radial and axial diffusivity and intracellular volume fraction (Cox et al., 2016), cortical measures such as regional gyrification (Gregory et al., 2016) and grey matter density (Ruigrok et al., 2014), and pathological brain structures such as white matter hyperintensities (Wardlaw et al., 2015) and enlarged perivascular spaces (Potter et al., 2015) may show interesting patterns of sex differences both across the population, and in how they relate to healthy behavioural variation as well as disease states.

### Conclusions

The present study is the largest single-sample study of neuroanatomical sex differences to date. We report evidence on the pattern of sex differences in brain volume, surface area, cortical thickness, white matter microstructure, and functional connectivity between adult males and females in the range between middle - and older-age. As has previously been argued (Fine, 2017), providing a clear characterisation of neurobiological sex differences is a step towards understanding patterns of differential prevalence in neurodevelopmental disorders such as autism spectrum disorder (Baron-Cohen et al., 2011), a variety of psychiatric conditions such as schizophrenia (Aleman et al., 2003), and neurodegenerative disorders such as Alzheimer’s Disease (Mazure & Swendsen, 2016; Viña & Lloret, 2010). We hope that the results provided here, given their large-scale, multimodal nature, will constitute an authoritative point of reference for future studies on a wide range of questions on brain sex differences. Insights into how and where the brain differs as a function of sex— with considerably more precision than in previous investigations—will enable more targeted examinations into potential drivers of these differences across psychiatric, psychological, and other domains. In particular, integrating macrostructural, microstructural, and functional data is an important long-term goal (Gur & Gur, 2017). Data on many thousands of further MRI scans (to a maximum sample of 100,000 with MRI data) will be available from UK Biobank in the coming years, in addition to more complex cognitive testing batteries and genotypic data. Future studies will be able to explore in much greater depth the links between sex differences in the brain, their possible causes, and their potential medical and behavioural consequences.

## Acknowledgements

We are grateful to the UK Biobank members for their participation in the study, and to the UK Biobank team, who collected, processed, and made available the data for analysis. This work was carried out under UK Biobank application number 10279. The work was primarily carried out in The University of Edinburgh Centre for Cognitive Ageing and Cognitive Epidemiology (CCACE), is part of the UK Research Councils’ cross-council Lifelong Health and Wellbeing initiative (MR/K028992/1). Additional funding has been received from the Biotechnology and Biological Sciences Research Council (BBSRC), the Medical Research Council (MRC), and Age UK’s Disconnected Mind project. Authors S.J.R., S.R.C., D.C.M.L., and I.J.D. were supported by the MRC award to CCACE (MR/K026992/1). Author S.R.C. was supported by MRC grant MR/M013111/1. Author I.J.D. is additionally supported by the Dementias Platform UK (MR/L015382/1). The work was also supported by a Wellcome Trust Strategic Award, “Stratifying Resilience and Depression Longitudinally” (STRADL; reference 104036/Z/14/Z). Author X.S. receives support from the China Scholarship Council. Author L.M.R. was supported by an Erasmus Traineeship Grant. Author H.C.W. is supported by a JMAS SIM fellowship from the Royal College of Physicians of Edinburgh, and by an ESAT College Fellowship from the University of Edinburgh. Authors A.M.M., H.C.W., and S.M.L. gratefully acknowledge the support of the Sackler Foundation. Author A.M.M. has previously received grant support from Pfizer, Lilly, and Janssen for work completely unrelated to the data or analyses in this study. We are grateful to Anne Scheel, Gina Rippon, Michel Nivard, and Odile Fillod for helpful comments on a previous draft. None of the funders or other acknowledged individuals are responsible for our analysis or interpretation of our results.

